# Towards end-to-end disease prediction from raw metagenomic data

**DOI:** 10.1101/2020.10.29.360297

**Authors:** Maxence Queyrel, Edi Prifti, Alexandre Templier, Jean-Daniel Zucker

## Abstract

Analysis of the human microbiome using metagenomic sequencing data has demonstrated high ability in discriminating various human diseases. Raw metagenomic sequencing data require multiple complex and computationally heavy bioinformatics steps prior to data analysis. Such data contain millions of short sequences read from the fragmented DNA sequences and are stored as fastq files. Conventional processing pipelines consist in multiple steps including quality control, filtering, alignment of sequences against genomic catalogs (genes, species, taxonomic levels, functional pathways, etc.). These pipelines are complex to use, time consuming and rely on a large number of parameters that often provide variability and impact the estimation of the microbiome elements.

Training Deep Neural Networks directly from raw sequencing data is a promising approach to bypass some of the challenges associated with mainstream bioinformatics pipelines. Most of these methods use the concept of word and sentence embeddings that create a meaningful and numerical representation of DNA sequences, while extracting features and reducing the dimensionality of the data. In this paper we present an end-to-end approach that classifies patients into disease groups directly from raw metagenomic reads: *metagenome2vec*. This approach is composed of four steps (i) generating a vocabulary of k-mers and learning their numerical embeddings; (ii) learning DNA sequence (read) embeddings; (iii) identifying the genome from which the sequence is most likely to come and (iv) training a multiple instance learning classifier which predicts the phenotype based on the vector representation of the raw data. An attention mechanism is applied in the network so that the model can be interpreted, assigning a weight to the influence of the prediction for each genome. Using two public real-life data-sets as well a simulated one, we demonstrated that this original approach reaches high performance, comparable with the state-of-the-art methods applied directly on processed data though mainstream bioinformatics workflows. These results are encouraging for this proof of concept work. We believe that with further dedication, the DNN models have the potential to surpass mainstream bioinformatics workflows in disease classification tasks.

## 1 Introduction

Recent technological advances made it possible over the past decade to collect DNA from nearly all accessible ecosystems and sequence it with extremely high resolution at an increasing low cost. Such developments made it possible for a whole new field - that of metagenomics, to develop very rapidly (*Oulas* et al. 2015; *Mardis* 2017). Although in theory it can be used all kinds of organisms, metagenomics is mostly used to explore the structure of microbial communities living in a given ecosystem. The microbiome was shown to play a crucial role not only in the environmental ecosystems but also in relation to the host when they inhabit it. This is also the case for humans. When the gut microbiome is altered, which is also the largest reservoir of bacteria inhabiting humans and reaching up to several kilograms, it is often reflected as an alteration of the human health. Indeed, recent abundant research has demonstrated the strong relationship between these microorganisms and complex and chronic human diseases such as diabetes, cirrhosis, autism, obesity and more (D. *Liang* et al. 2018). The field is maturing rapidly and large public repositories store an increasing number of standardized datasets. Data collection is performed using Next Generation Sequencing (NGS), taking advantage of parallelization of DNA sequencing (shotgun), which generates millions of sequence reads in a short time. Reads are short DNA sequences (generally between 50 to several thousands of nucleotides (A, C, G, T) and are stored in standardized files such as the fastq. The development of metagenomic sequencing came along with the rapid development of bioinformatics workflows, which ultimately yield quantitative measurements of biological objects such as genes, species, genera and other taxonomic levels, functional pathways, etc in the form of count tables (*Nayfach* et al. 2016; *Wen* et al. 2017). Several steps are required to obtain such count tables and all of them rely on assumptions that affect the final outcome. The complex bioinformatics workflow starts by reading the fastq files and using the quality scores to filter out nucleotides as well as reads that do not pass the decided confidence criteria. Next, the reads are aligned onto the host genome to filter them out, while focusing on the microbiome data. Finally, the resulting reads are grouped together (binned) using different techniques, either through alignment with reference gene/genome catalogues or thorough other approaches based on k-mer similarity or co-abundance clustering (*MetaHIT Consortium*, *Nielsen*, et al. 2014, *Qian* et al. 2019). After these bioinformatics processing steps, the data needs to be prepared for further statistical analyses including constructing machine learning models.

The bioinformatics workflows are dependent on different software, which are not designed to work together in the most efficient way. Moreover, a large number of parameters and thresholds are to be set, often arbitrarily that affect the final outcome. It is however possible in the context of microbiome classifications, to follow another strategy and reduce the complexity of the bioinformatics workflows described above. In recent years, distributed representations of words in a vector space has been increasingly used in Natural Language Processing (NLP) to improve the performance of learning algorithms (*Mikolov* et al. 2013). These representations, called embeddings, are projection of words in a numerical vector space capturing semantic and lexical information learned with contexts of words. These vectors can be used as features for many applications like sentiment analysis (*Maas* et al. 2011), translation (*Qi* et al. 2018) or even speech recognition (*Bengio* et al. 2014), outperforming standard word count representation. In metagenomics, DNA sequences can be embedded in a similar way with some pre-processing (*Ng* 2017; *Menegaux* et al. 2019; *Joulin* et al. 2016). A DNA sequence is composed of four nucleotides A, C, G and T. Consequently, it may be similar to a sentence in NLP with a shorter alphabet. However, it is necessary to take into consideration other distinctions between metagenomes and NLP data. DNA sequences do not have concept of word delimited by spaces between letters. To bypass this, DNA sub-strings called k-mers (Figure 1) may be introduced. K-mers are composed of *k* nucleotides and refer to all sub-sequences from a read obtained by DNA sequencing. The possible amount of k-mers given a read of length L is *L* − *k* + 1, and the possible number of combinations is equal to 4^*k*^ (since there are 4 distinct nucleotides). Furthermore, NGS makes “massively parallel sequencing” to numerically convert several DNA fragments from different sources into short reads. Thus, metagenomic data is composed of several sequences with any information about the order. This is very different since textual data know order between sentences or paragraphs.

**Figure 1:**
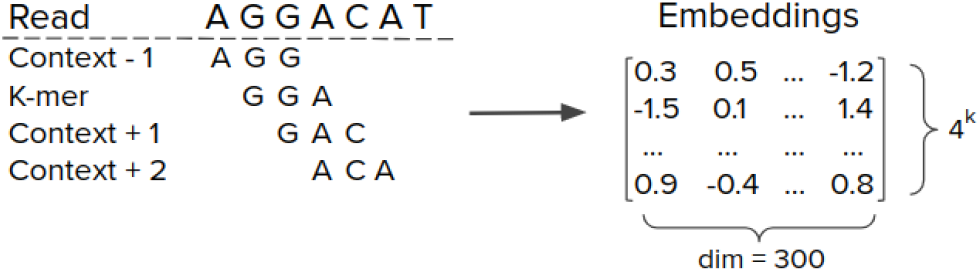
Left side represents a read cut into k-mers of length 3 with a window size of 2 and a padding size of 1. Right side corresponds to an embeddings matrix of dimension 300 learnt with k-mers vocabulary of size 4^*k*^ = 4^3^ = 64.

In this proof of concept paper, we documented an implementation of the idea that deep neuronal networks can bypass the classical bioinformatics workflow and allow to classify metagenomic samples directly from raw sequence data. Moreover, the trained DNN may automatically discover important biological concepts that could be further used to understand the contextual signatures of the classification process. Besides the robustness that this approach might offer, it could also prove to be computationally more efficient. Indeed, to predict the class of one sample, the time required in bioinformatics workflows is between 1 to 3 hours (*Ugarte* et al. 2019) whereas this method requires less than one hour. Such an end-to-end approach could be very useful in the clinical setting, without needing to send out the data for heavy processing to bioinformatics platforms.

The core of our approach lies on the integration of different types of embeddings that encode the metagenomic sequences. We divide this approach into four main stages and assign a name to each of them for more clarity. The first one, *kmer2vec*, consists of a transformation of k-mers into embeddings. The second, *read2vec*, refers to read projection into embeddings. *kmer2vec* and *read2vec* acts in a similar way to NLP which transforms words and sentences into vectors. The third, *read2genome*, classifies reads onto bacterial genomes from which they most likely originated. The purpose of this step was to group similar read embeddings in order to amplify their information.

The fourth and last step, *metagenome2vec*, begins by transforming the metagenomes onto robust multiple instance representations using *read2vec* and *read2genome* and drastically reduce the initial dimensional complexity. Finally, a multiple instance learning (MIL) classifier is trained on the transformed metagenomes’ vector space using labels from the classes to discriminate.

## 2 The representation of metagenomic data with embeddings

### Representing Metagenomes

When using microbiome data in a precision medicine context, one critical task is to discriminate diseased patients from healthy or within different severity groups (*Jobin* 2018). To represent mathematically the different concepts of our approach, let 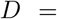 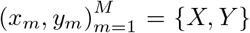 denote a set of *M* metagenomes and the associated labels *Y* ∈ {0, 1}^*M*^. A metagenome *x*_*m*_ is composed of *R*_*m*_ ≫ 10^6^ DNA reads. A DNA read *s*_*r*_, *r* ∈ {0..*R*_*m*_} is a sequence of length *L*_*r*_ ∈ 50 ~ 200 formed by several nucleotides in the vocabulary 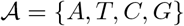, so 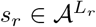.

### Representing reads

In NLP, there are explicit word and sentence delimiters. On the contrary, in the case of DNA reads there is no explicit information to systematically delimit sub-sequences. Moreover, it is difficult to know the location of a read in genomes because the DNA were fragmented prior to sequencing and there is no particular order to the reads after sequencing. To transform the reads onto something similar to words, a possible approach may be to simply split the sequences into k-mers (*Menegaux* et al. 2019; *Woloszynek* et al. 2018; *Min* et al. 2017; Q. *Liang* et al. 2020). Various size of *k* can be considered depending on the task. Padding between k-mers is equal to one because k-mers refers to all sub-sequences of length k. For example, if we have a sequence of seven bases “AATCCGA” and if *k* = 3, then after splitting we do not only obtain the k-mers “AAT” and “CCG” but also “ATC”, “TCC” and “CGA”.

### Building k-mers, reads and metagenome embeddings

Similar to the NLP applications, using vector representations of k-mers can overcome some limitation to treat reads as discrete atomic sequence of nucleotides. Similarly, vector representation of reads and metagenomes can be envisioned to go beyond their simple encoding representations (*Woloszynek* et al. 2018). In this paper we focus on learning metagenome embeddings that have the ability to both reduce the data dimensions and also support computationally-efficient predictive models from embedded raw metagenomes. As Metagenomes are composed of reads and reads are composed of k-mers, it is natural to consider a multilevel embeddings approach. This is the reason why we have introduced three main stages of data transformation (Section 1): *kmer2vec*, *read2vec* and *metagenome2vec* to compute respectively vector representations of k-mers, reads and metagenomes. These transformations are detailed in Section 4.

## 3 Related work

Several recent deep learning methods have been adapted to support the analysis of genomic or metagenomic data for different tasks. Recall that genomics differs from metagenomics in the fact that in the latter multiple organisms are considered in the same time and their DNA is mixed together and consequently the sequenced data as well. So the differences between these studies are the representation of DNA sequences, the types of algorithms and obviously the final objective. The Figure 3 in appendix summarizes the different metagenomic embeddings models in relation to our approach *metagenome2vec*.

### Machine learning models from nucleotide one hot encoding

It is a well known that recurrent neural network (RNN) as well as convolutional neural networks (CNN) both work well on sequential data (*Cui* et al. 2016; *Young* et al. 2018). That is why, Genomic studies have used both architectures on DNA sequences. One classical genomic task is to analyze *chromatin accessibility*. It involves in finding some regions of the genome that are accessible for cellular machines involved in gene expression while others are compactly wrapped and remain inaccessible. Different works have shown that RNN or CNN on DNA sequences can capture relevant patterns and outperform state-of-the-art methods in chromatin accessibility prediction for instance (*DanQ* - (*Quang* et al. 2016), *DeepSEA* - (*Zhou* et al. 2015), *Basset* - (*Kelley* et al. 2016)). A benchmark of these approaches was introduced by *Hildick-Smith* et al. (2016).

In metagenomics, some studies focus on the hierarchical taxonomy structure and are faced with a multi-class classification problem. This is the case of *GeNet* (*Rojas-Carulla* et al. 2019), a deep learning model based on CNN and ResNet architecture (*He* et al. 2015). Authors use a one-hot encoding of the input nucleotides and their position in the read. The loss function is computed at each taxonomy level and the prediction at any level is forwarded to the next one adjusting the decision of the classifier.

However, all these algorithms keep the initial representation of DNA and simply one-hot encode the four nucleotide bases {A, C, G, T}. In other words, most algorithms operate on a 4 × *L*_*r*_ matrix where *L*_*r*_ is the sequence length. This representation is quite basic and does not consider the similarities between k-mers. This is comparable to representing a sentence as a set of letters rather than a set of words.

### Machine learning models from k-mers embeddings

Research in NLP has seen a major development on low-dimensional representation of words. These methods regularly outperform the simple version of bag of words by projecting words into a vector representation that accurately captures syntactic and semantic word relationships. Recently, based on this concept, there has been some approaches considering k-mers embeddings. In the work of *Ng* (2017), k-mer embeddings are computed with a *word2vec* algorithm. A relationship is highlighted between the arithmetic operation on *word2vec* vectors and the corresponding concatenation of k-mers. Similarly to the study *Min* et al. (2017), where the goal is to classify chromatin accessibility (as we saw in the previous paragraph), the *GloVe* algorithm (*Pennington* et al. 2014) is used to create k-mer embeddings before training the final neural network. Experiments have shown that results are better when the sequence is transformed into k-mer embeddings. Nevertheless, in their study *Glove* set the k-mer size to 6 without discussing other configurations that could have demonstrated the overall importance of this parameter.

There are also several attempts to learn machine learning models directly from raw metagenomic data. Most of them address the task of predicting the origin of reads (called taxonomic profiling) at a certain taxonomy level or to perform phenotype classification. To assign taxonomic information to each read, the authors of *FastDNA* algorithm (*Menegaux* et al. 2019) have demonstrated that their approach using embeddings of k-mers achieves performances comparable to the state-of-the-art. In the first step of their approach they define the length k of the k-mers that describe the DNA sequences. Then they run the *fastText* algorithm (*Joulin* et al. 2016) to learn low-dimensional embeddings (dimension from 10 up to 1000). All k-mers in a sequence are replaced by their corresponding vector and summed to get an embedding of the read they belong to. Then this new vector is passed to a linear classifier, which computes the softmax function and minimizes the cross-entropy loss by gradient descent. The k-mer embeddings are directly learned from the read classification considering the result of the loss function. The authors demonstrated that significant results of prediction appear with k-mer size above 9 nucleotides. With a similar objective, Q. *Liang* et al. (2020) propose *DeepMicrobes*, a neural network with an architecture composed of an embedding of k-mers, a bidirectional LSTM, and a self-attention layer to predict the species or the genus of a read. In their experiments, k-mers of size k=12, lead to their best results.

In the work of *Woloszynek* et al. (2018), the objective is to add, in addition to taxonomic profiling, a method to retrieve the source environment of a metagenome (phenotype prediction). A *Skip-gram word2vec* algorithm (*Mikolov* et al. 2013) is trained for k-mers embeddings and a SIF algorithm (*Arora* et al. 2017) is used to create reads and samples embeddings. They demonstrate the usefulness of such an approach for clustering and classification tasks. Moreover, they show that such embeddings allow models to be trainable using k-mers with big *k* (greater than 9), which is not possible when relying on simpler representation such as one-hot encoding because of their size exponentially growing with k.

## 4 *metagenome2vec*: an approach to learn metagenomes embeddings

We introduce *metagenome2vec*, a method to transform shotgun metagenomic data into a suitable embedding representation for downstream task such as disease classification. *metagenome2vec* is trained from raw DNA sequences through several specific steps. Indeed, metagenome embeddings are built from embeddings of reads themselves built from k-mers embeddings. We highlighted state-of-the-art algorithms that learn embeddings of k-mers and reads. The proposed approach is based on multiple instance learning to construct metagenome embeddings. The global architecture is summarized in Appendix B and all blocks of the pipeline are explained below.

### 4.1 kmer2vec: learning k-mers embeddings

DNA sequences are split into several k-mers before learning k-mers embeddings. The context of a k-mer corresponds to both the preceding and the following k-mers. The context can consist of several k-mers; this parameter is called the window size and tuned to the number of surrounding context k-mers desired. After learning, all k-mers are indexed in the embeddings matrix. Figure 1 illustrates this process.

### 4.2 read2vec: Learning Read Embeddings

It has been shown that the algorithms constructing word embeddings give good results for representing short sentences by simple arithmetic operations on word vectors (*Mikolov* et al. 2013; *Pennington* et al. 2014; *Joulin* et al. 2016). Nevertheless, more sophisticated approaches for sentence embeddings have been developed. Sequence-to-sequence models (*Sutskever* et al. 2014) for instance, use a first network called the *encoder* to encode the sentence information. And a second one, called *decoder* decodes the sentence information for a specific task such as a translation where the authors have demonstrated great performance. The hidden layers of the encoder represent the sentence embeddings. *Skip-thought* (*Kiros* et al. 2015) vectorizes sentences with this approach, learning to generate surrounding sentences. *SDAE* (*Hill* et al. 2016), uses a LSTM encoder-decoder to reconstruct noisy sentences (words are removed or switched places according to a probability). More recently, *BERT* model (*Devlin* et al. 2018) trains an encoder with self-attention mechanism (*Luong* et al. 2015), called transformer, to learn contextual relations between words to retrieve masked words or predict the next sentence. DNA sequences can be transformed into embeddings using previous algorithms in the same way as sentences are.

It is also possible to train a supervised model on sequences of letters. In NLP, learning a supervised model is generally related to language inference, which is the task of determining whether a “hypothesis” is true (implication), false (contradiction), or indeterminate (neutral) given a “premise” (*InferSent* algorithm (*Conneau* et al. 2017) works with this concept). Since there is no information about textual concepts to learn about DNA sequences, another approach is to train the model by predicting the taxonomy of a read as it is performed with *FastDNA* (Section 3).

In our experiments, three algorithms were integrated in the workflow. *fastText* (see Appendix C for more details) sentence embeddings as implemented in the package, *FastDNA* sequence embeddings extracted after the model were trained to classify taxonomy reads at the species level and finally a sequence-to-sequence based transformer applied on language modeling task (determine the probability distribution for the likelihood of a given word to follow a sequence of words). Each of these algorithms respect two properties: (i) being efficient enough to process the millions of sequences per metagenome (a non-blocking point in theory but important for implementation), and (ii) not involving sentence order in the prediction task.

### 4.3 read2genome: reads classification

We would like to take advantage of the putative origin of the reads to construct metagenome representation. *read2genome* then acts as a clustering process producing bag of reads with genome similarity. To address the question of predicting the genome to whom a read most probably belongs, we have relied on two methods in our experiments. Firstly, *FastDNA* (*Menegaux* et al. 2019) that learns embeddings and classification with an end-to-end supervised model. Secondly, a Multi Layer Perceptron (MLP) classifier that takes as inputs the read embeddings learnt by self-supervised training with *read2vec*. Appendix G highlights evaluation metrics used for *read2genome*.

### 4.4 metagenome2vec: learning metagenome embeddings

The following step is to create metagenome embeddings using read embeddings or a set of reads embeddings. We propose to consider two different approaches in building metagenome embeddings: (i) the vectorial representation as baseline and (ii) the multiple instances learning representation as our reference method. The notations in the next sections are in accordance with those introduced in section 2.

#### 4.4.1 metagenome2vec: Vectorial representation

Once a low-dimensional representation of the reads is available, all reads from a given metagenome are transformed into embeddings and summed together, resulting into a single instance embeddings for one metagenome. A representation of metagenome can be computed as shown in the equation 1:

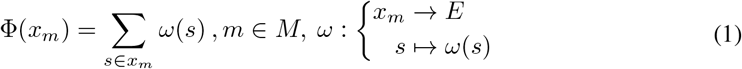

With *ω* the *read2vec* transformation, *x*_*m*_ the ensemble of reads in the metagenome *m* and *E* the dimension of the embeddings.

The tabular representation is the one used by most machine learning algorithms. It relies on a more abstract representation than the multiple instances representation. Indeed, all the information of an entire metagenome, its millions of reads related to hundreds of different genomes, is summarized into a unique vector in the latent embeddings space. In addition to the evaluation of the classification we have defined an analysis based on clustering detailed in the Appendix H.

#### 4.4.2 metagenome2vec: Multiple instances learning representation

Metagenomic data can be thought as a set that contains millions of reads representing one sample. The size of a set can varies depending on the sequencing technology and there is no specific order between reads within a set. Learning from such bags of reads correspond to a Multiple-Instance Learning (MIL) problem that deals with an objective function that is invariant to permutation and operates on non-finite dimensional vectors. In this study, we implement a deep learning algorithm from the work of *Zaheer* et al. (2017) named *DeepSets* and an extension of the multiple instance layer with an attention mechanism from *Ilse* et al. (2018) (see Appendix D for more details about these algorithms). The attention mechanism assigns a weight for each instance to determine which one in the set helped to predict the label. As the metagenomes are represented as bag of genomes embeddings, it is interesting to integrate such an aggregation operation to determine the taxa that play a bigger role in the prediction. In that way the model could be interpretable. Model interpretability is essential for the trust of application with predictive models for metagenomics-based precision medicine, as it would provide usable information for precision medicine initiatives (*Prifti* et al. 2020).

Instead of aggregating all the computed embeddings reads to form one vector, a first idea is to keep this representation to save all information. Unfortunately, one metagenome is composed to potentially millions of reads. Thus, a bag is far too large to fit in memory for further processing by the machine learning algorithms. We support another approach, consisting of first training a classifier (*read2genome*) to predict the genomes from which the DNA sequences may have originated.

Rather than summing all read embeddings as in the previous method, it is possible to sum embeddings of reads belonging to the same taxonomic levels, namely genome or genus. As a result, each metagenome is represented by a set of embeddings. In the literature learning from objects described as sets of tabular value is called a MIL problem (*Wang* et al. 2000) where a class label is assigned to a bag of instances. Figure 2 schematizes the model and the equation 2 that presents these concepts is described below:

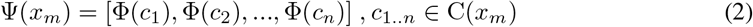

Where C is the *read2genome* function clustering reads of a metagenome into *n* clusters, thus *c*_*n*_ is a group of reads, *x*_*m*_ the ensemble of reads in the metagenome *m* and *c*_*n*_ ⊂ *x*_*m*_.

**Figure 2:**
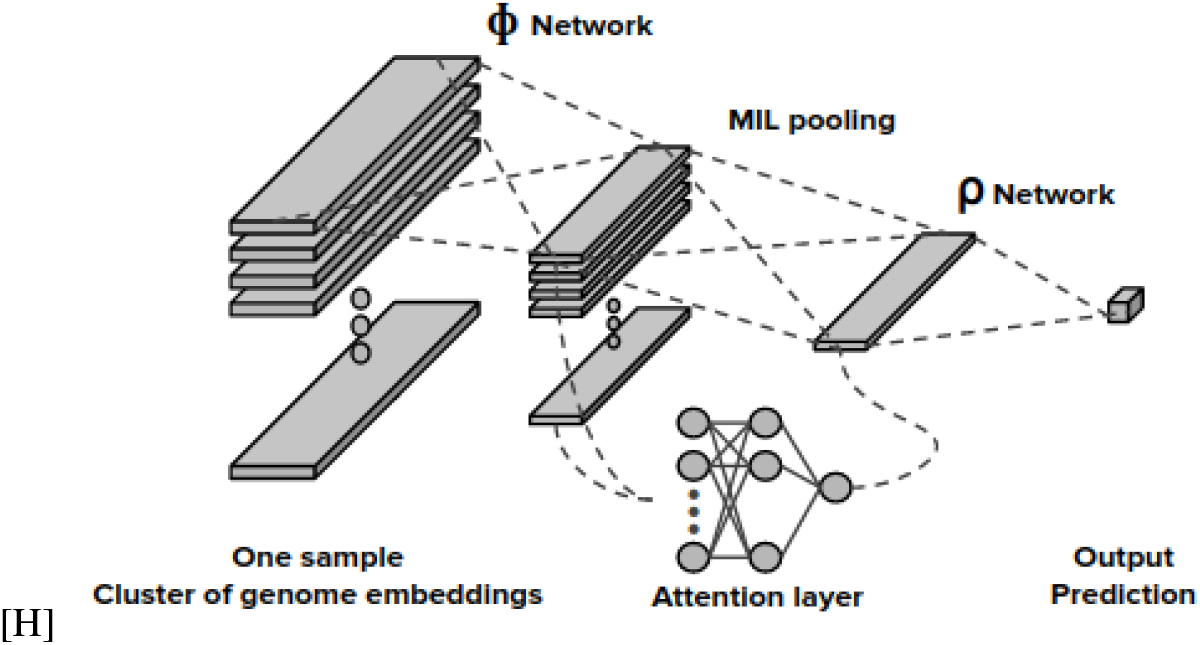
DeepSets neural network architecture with attention as MIL layer. The input is a set of genome cluster embeddings and the output is the phenotype prediction.

As reads are fragments of DNA from several genomes, grouping them into clusters projected onto the embedding vector space could bring specific information for each metagenome.

## 5 Experiments and Results

We devised several experiments to test the efficacy and the efficiency of our novel *metagenome2vec* algorithm to classify metagenomes onto classes of diseases with which the hosts are associated. The performance of *metagenome2vec* w.r.t. the state-of-the-art was tested on two benchmark disease classification tasks (predicting colorectal cancer and liver cirrhosis (*Edoardo Pasolli* et al. 2015)) as well as a simulated data-set). Moreover, to understand the source of power of the *metagenome2vec* algorithm, we also tested the intrinsic quality of the learnt embeddings and the ability to assign a read to the right taxa.

### 5.1 Datasets

#### Datasets used to learn embeddings: *kmer2vec* and *read2vec*

The catalog we used here was composed of 506 different genomes, that were used to build the IGC gene catalog (*MetaHIT Consortium*, *Li*, et al. 2014). These genomes correspond to 235 species, 79 genera and 37 families.

#### Datasets to learn Read2Genomes classifier

The *CAMISIM* (*Fritz* et al. 2019) software was used to simulate 5,11M and 2,19M DNA sequences (for the train and validation set respectively) based on the genome catalog mentioned above. This allowed us to train and test the DNN models in how well their performed in estimating the abundance profiles resulting from read classification onto the genomes. The reference abundance used for the simulation followed a uniform distribution so that it would be balanced with a number of sequences almost equal for each genomes. This is not the case in real metagenomes where abundance follows exponential distributions. Nevertheless, in the case of read classification modeling, this avoids the classifier to focus and predict the predominant classes, thus learning more robust embedding.

#### Datasets to learn the Disease prediction tasks

Our experiments were performed on two datasets with controlled disease liver cirrhosis (Cirrhosis) and colorectal cancer (Colorectal) from UCBI open source studies. In colorectal datast 15 patients had an adenoma, this is a benign tumor so they have been labeled as control cases as in *Pasolli* et al. (2016). We also artificially simulated a third dataset (Simulated) with an artificial disease using *CAMISIM* (*Fritz* et al. 2019) generator based on known bacterial genomes and abundance profiles which significantly vary in abundance between two groups (namely disease vs not). The control case samples are simulated with the same abundance of genomes, thus the number of reads is almost the same for each species. In contrast, the samples from diseased patients had an abundance 3 times more important for five species that imply the disease. Datasets have a massive size considering they only have few hundreds of patients. Metagenomes are composed of ~ 80 million reads, each one composed of ~ 90 nucleotides. It is a case where *D* >> *N* with *D* the number of features and *N* the number of examples. This is, in general, a complex task for machine learning algorithms. Information about datasets are described in Table 1.

**Table 1:** Information about the metagenomic datasets

| Dataset name | Disease | Control subjects | Case subjects | Control-to-Case | Size<sup>1</sup> | Reference |
| --- | --- | --- | --- | --- | --- | --- |
| Colorectal | Colorectal Cancer | 73 | 48 | 60.3% | $\sim 866Go$ | Zeller et al. 2014 |
| Cirrhosis | Liver Cirrhosis | 114 | 118 | 49.1% | $\sim 1To$ | Qin et al. 2014 |
| Simulated | Artificial | 100 | 100 | 50.0% | $\sim 10Go$ | - |

### 5.2 Reference Methods compared to metagenome2Vec

To our knowledge there is no other study applying machine learning directly on raw metagenomic data to predict disease. In general, disease classification with metagenomic data is done with standard pipeline using species-level relative abundances and presence of strain-specific markers (*Pasolli* et al. 2016). On top of these bioinformatic processes, machine learning algorithms like SVM, Random Forest or Elastic Net are trained to make predictions. More recently, *Oh* et al. (2020) have proposed to highlight the use of auto-encoder models on such metagenomic abundance tables. Results are reported in the table 2 and are used in this paper as part of the state-of-the-art benchmark.

**Table 2:** Classification metrics for three databases Colorectal, Cirrhosis and Simulated. The standard deviation is calculated for the accuracy score and written in parenthesis. Results are reported for two reference methods (*MetaML* and *DeepMicro*) that use species-level relative abundances and presence of strain-specific markers. *BoK*, *M2V-VR* and *M2V-MIL* are methods tested in this study. For each method, all the scores are the best of all the experiments.

| Method | Metrics | Colorectal | Cirrhosis | Simulated |
| --- | --- | --- | --- | --- |
| MetaML | Accuracy | 0.81<br>(0.068) | 0.88<br>(0.043) | - |
|  | precision | 0.82 | 0.89 | - |
|  | recall | 0.81 | 0.88 | - |
|  | F1-score | 0.79 | 0.88 | - |
|  | AUC | 0.87 | 0.95 | - |
| DeepMicro | AUC | 0.81 | 0.94 | - |
| BoK<br>k=3 / k=6 / k=9 | Accuracy | 0.64 / 0.69 / 0.65<br>(0.061) / (0.055) / (0.071) | 0.67 / 0.73 / 0.75<br>(0.077) / (0.062) / (0.043) | 0.61 / 0.66 / 0.63<br>(0.032) / (0.040) / (0.057) |
|  | precision | 0.81 / 0.88 / 0.84 | 0.66 / 0.76 / 0.77 | 0.61 / 0.66 / 0.71 |
|  | recall | 0.57 / 0.61 / 0.60 | 0.77 / 0.69 / 0.74 | 0.56 / 0.62 / 0.55 |
|  | F1-score | 0.55 / 0.59 / 0.56 | 0.70 / 0.72 / 0.74 | 0.59 / 0.64 / 0.65 |
|  | AUC | 0.66 / 0.73 / 0.67 | 0.71 / 0.77 / 0.77 | 0.65 / 0.69 / 0.67 |
| M2V-VR | Accuracy | 0.72<br>(0.058) | 0.79<br>(0.044) | 0.85<br>(0.047) |
|  | precision | 0.77 | 0.78 | 0.87 |
|  | recall | 0.64 | 0.67 | 0.81 |
|  | F1-score | 0.69 | 0.70 | 0.84 |
|  | AUC | 0.794 | 0.81 | 0.85 |
| M2V-MIL | Accuracy | 0.81<br>(0.065) | 0.82<br>(0.056) | 0.92<br>(0.070) |
|  | precision | 0.78 | 0.83 | 0.94 |
|  | recall | 0.76 | 0.84 | 0.91 |
|  | F1-score | 0.76 | 0.83 | 0.92 |
|  | AUC | 0.81 | 0.83 | 0.92 |

### 5.3 Results of the Disease prediction tasks

All experiments are done by limiting to one Million the number of reads for each metagenome. Indeed, learning computation time can take several days and there are many parameters to test for both structuring and machine learning. This is a way to cover many experiments and save time before running on the whole data. The datasets are composed by only hundreds of samples so to tune the hyper parameters and avoid over fitting we apply a nested cross validation. In that way, 20% of the data form the test set, the 80% remaining are used to perform a 10-fold cross validation to tune hyper parameters with 20% of the data as validation set. The whole operation is repeated 10 times with different train and test sets. The tuning is driven by accuracy score. AUC, precision, recall and F1 score are also computed. Details about code implementation are given in appendix E.

*MetaML* (*Pasolli* et al. 2016) and *DeepMicro* (*Oh* et al. 2020) are the reference methods. Bag of K-mers (*BoK*) method is related to Bag of Word (*BoW*). Thus, *BoK* represents a metagenome as a vector by counting all the occurrences of its k-mers without using embeddings. is the baseline to confirm embeddings usefulness. M2V-VR is “*metagenome2vec* vectorial representation” and M2V-MIL is “*metagenome2vec* multiple instance learning representation”. Colorectal and Cirrhosis datasets are benchmarked with all previous methods whereas the Simulated dataset is used to evaluate M2V-MIL efficiency and interpretability. Models used for BoK and M2V-VR methods are tuned with random search using 100 different sets of parameters. M2V-MIL is tuned with approaches more adapted for model composed by a lot of hyper parameters like neural network. Thus, Bayesian optimization is applied for continuous parameters and bandit optimization for discrete parameters. The table 2 summarize our results.

The BoK baseline has been computed with three different values of *k* equal to 3, 6 and 9. Better results are obtained when *k* = 6 or *k* = 9 without a significant difference among them. It cannot be computed with higher value of *k* due to the number of distinct k-mers in the vocabulary that becomes too large. Scores are lower than other methods as expected but still leads quite good results for a simple, training free representation. Our results demonstrate that adding embeddings abstraction increases the performance and that the MIL representation yields better results. Our approach *M2V-MIL* reaches the same level as the state-of-the-art for Colorectal but is still lag behind for the Cirrhosis dataset. On the Simulated dataset, *M2V_MIL* provides better results. This dataset is useful to assess whether the MIL model retrieved the genomes that play a role in the classification or not. We extract, for each well-ranked positive (artificial disease) sample, the five main genomes that had the greatest impact on the outcome, resulting in a total of 12 distinct genomes. Among them, two of the five genomes invoking disease (altered abundance) were predicted at the species level at 18% and 14.5% on all well-ranked positive samples. The 12 genomes (from the top-5) are similar at the genus level with four of the genomes to be found. These results showed the descriptive accuracy of the models.

## 6 Conclusion

Phenotype prediction using metagenomic data is a critical task especially in a clinical setting. In this work, we proposed *metagenome2vec*, an algorithm which transforms raw metagenomic data onto automatically learned representations allowing us to apply on them multiple instance learning-based approaches. Combined with an attention mechanism, the algorithm offers some interpretability by being able to detect instances / genomes in the MIL set that could play a role in the phenotype prediction. Compared to standard quantitative metagenomics, our model displayed a performance comparable with the state of the art ML approaches directly applied to data processed through bioinformatics workflows. However, as it is an end-to-end algorithm, the time which would be required to run a costly bioinformatics pipeline is saved in inference mode. One of the limits of this work lies in the computational complexity of the learning of the different embeddings and the identification of each one’s contribution to the prediction power. That is why we have proposed some analyses for each step to facilitate their intrinsic evaluation and optimization. We plan to test a wider range of parameters in the pipeline along with various algorithms so as to read multi-purpose embeddings and test them on all existing public datasets related to other diseases (*Pasolli* et al. 2016).

## Appendix

## A Deep Learning workflow in metagenomics

**Figure 3:**
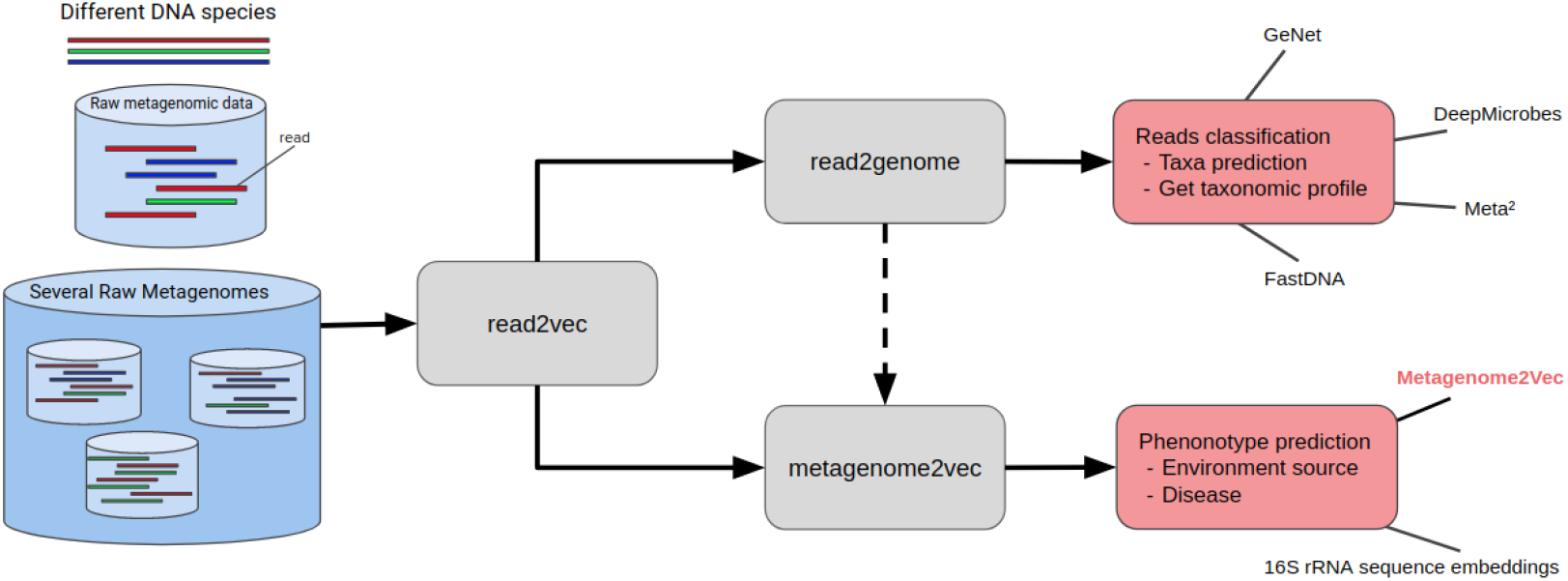
Work flow of metagenomic data projected into low-dimensional representation with embedding learning algorithms. The blue color represents the input data, the grey color represents different internal modules of the pipeline and the red color the prediction task performed. The dotted line is only a part of the *metagenome2vec* algorithm. Algorithms written (including *metagenome2vec*) are linked with their corresponding task. We can see that *read2vec* is a module for both phenotype and read classification. If the abstraction is at the read level, results are handled to classify reads for taxonomic profiling. If the abstraction is at the metagenome level, prediction could be used for phenotype classification

## B Architecture overview

In the following diagrams, the color blue corresponds to the metagenomic data, while yellow refers to the learned embeddings, and red represents the learning part of the module.

## B.1 Global architecture of metagenome2vec

**Figure 4:**
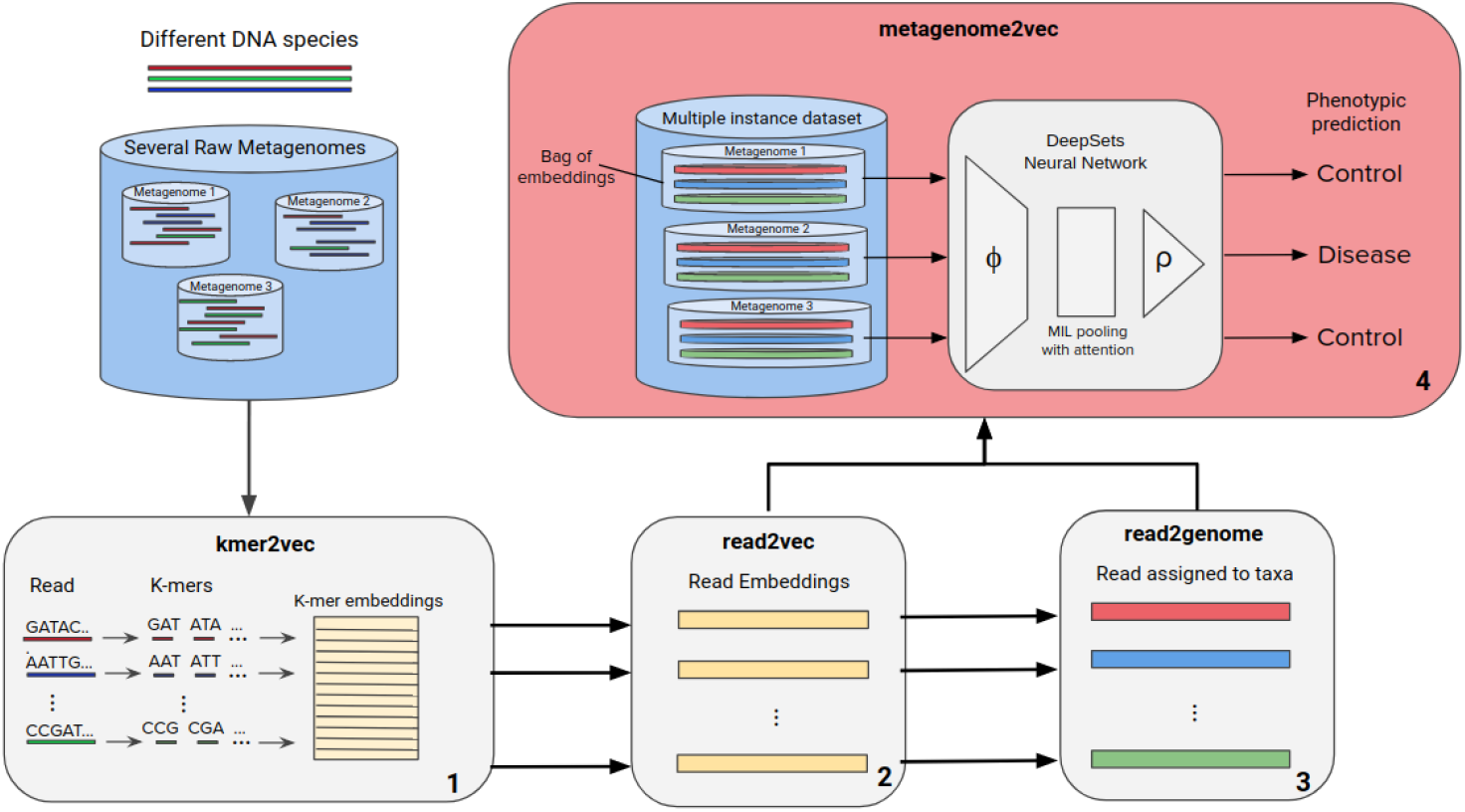
Numbers indicate the order of each operation block. Raw metagenomic data is the input of *metagenome2vec*. All DNA sequences are embedded by *kmer2vec* and *read2vec* algorithms (Figure 5 and 6). Then, *read2genome* (Figure 7) uses these embeddings to assign a cluster, corresponding to a genome id, for all reads. Embeddings of reads in the same cluster are averaged. It results in a multiple instance dataset where a bag of embeddings represents one metagenome. At the end, a multiple instance learning model DeepSets is trained to perform disease prediction.

## B.2 kmer2vec: Example with FastText

**Figure 5:**
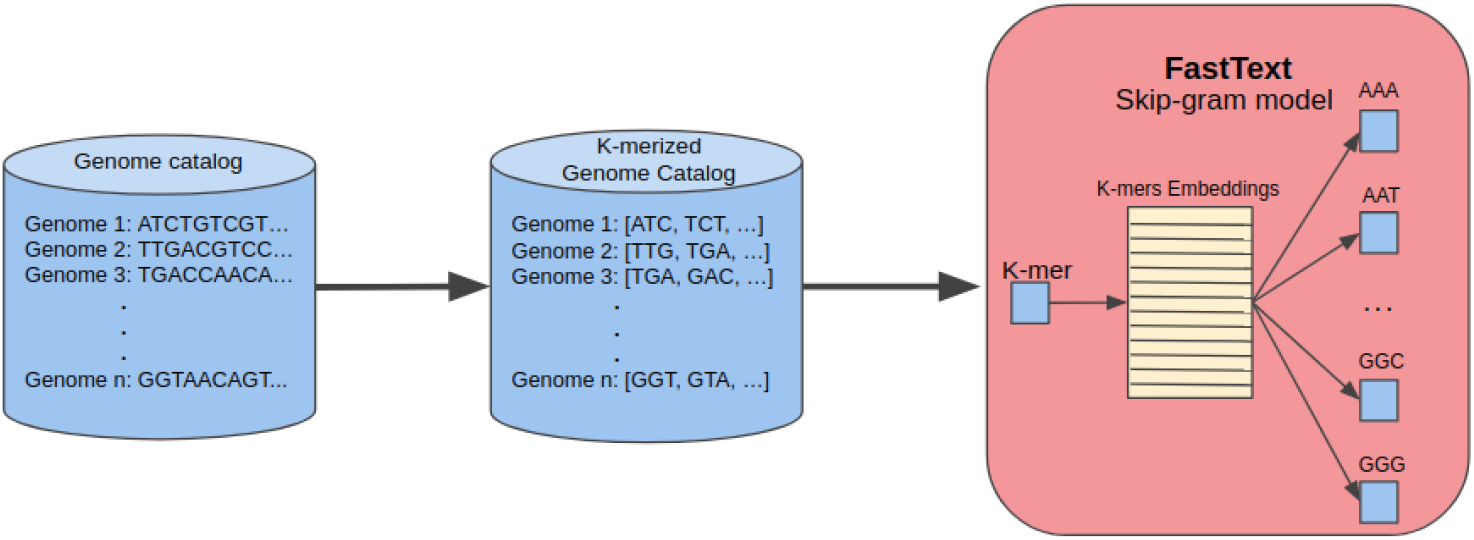
The first step is to transform genomes sequences into k-mers with a specific *k* size. On the figure k=3 because k-mers are composed of three nucleotides. Then, k-mers are passed to the *FastText skip-gram* model (Appendix C) learning to retrieve surroundings k-mers context.

## B.3 read2vec: Example with a transformer language model

**Figure 6:**
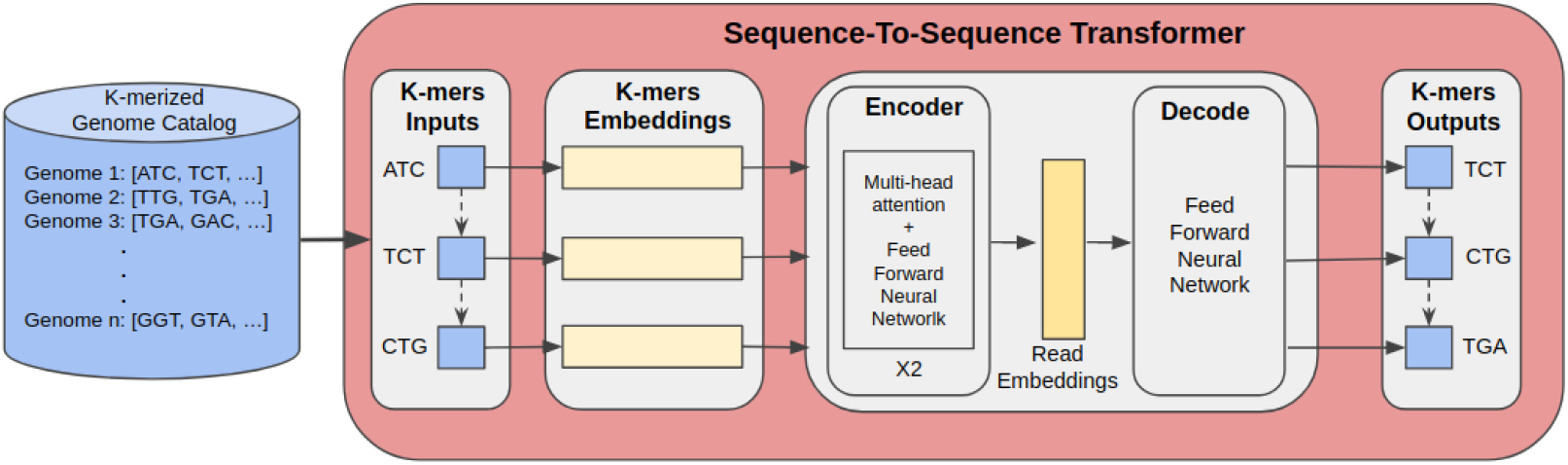
Sequences, cut into k-mers, pass into a transformer sequence-to-sequence language model. A first layer converts k-mers to their embeddings learnt in *kmer2vec* (Figure 5). The encoder creates the read embeddings with two blocks composed by a multi-head attention and a feed forwoard neural network. The decoder tries to predict the next k-mers from the source sequence passing the read embeddings in a fully connected layer before computing the softmax to get a probability for each k-mers. When *k* is relatively big, this last layer is quite intensive to compute because its complexity grows linearly with the size of the vocabulary. Thus, the adaptive softmax proposed by *Grave* et al. (2017) is used instead of softmax to be more efficient without reducing performance.

## B.4 read2genome

**Figure 7:**
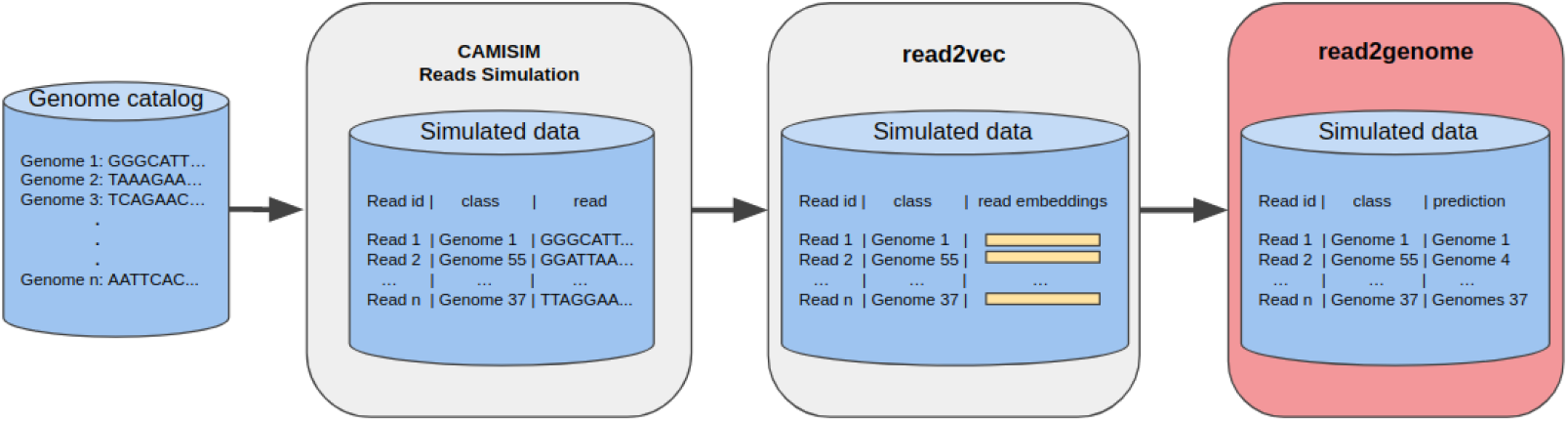
A catalog of complete genome is used by the *CAMISIM* software (*Fritz* et al. 2019) to simulate metagenomic data with a specific taxonomic profile (abundance of species). The resulting dataset is a set of reads associated with the identifier of the genome from which they originate. Reads are embedded by *read2vec* (Figure 6) before being passed into *read2genome* trained to retrieve their source genome.

## C FastText for k-mers embeddings

We first introduce the *word2vec skip-gram* model because FastText is a derivative of this algorithm.

## word2vec

This is the well known model popularized by Google with *Mikolov* et al. (2013). It is a shallow two-layer neural network auto-encoder. We opted for the *skip-gram* version of the model. So the neural network has been trained to predict the most obvious surrounding context for each k-mers. The prediction is based on the softmax function that gives the posterior distribution of k-mers:

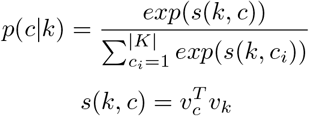

With *k* the current k-mer, *K* the vocabulary of k-mers, *c* the context k-mer, *v* represents a vector, *s*(*w, c*) denotes the scoring function between *w* and *c*.

## FastText

Published by Facebook (*Joulin* et al. 2016; *Bojanowski* et al. 2016), this algorithm brings the concept of *subword* information based on character n-grams. In fact, *fastText* works like the *word2vec skip-gram* model but with a different scoring function. In *word2vec* a dot product is done between the current word and its context words. Here, the dot product is computed between all corresponding characters from min-n-gram to max-n-gram (two hyper parameters). For example if we consider a k-mer “ATACCA”, min-n-gram=3 and max-n-gram=6, the n-grams are {ATA, TAC, ACC, CCA, ATAC, TACC, ACCA, ATACC, TACCA, ATACCA}. A k-mer is finally represented as the sum of the vector representations of its n-grams. The scoring function is re-written as follows:

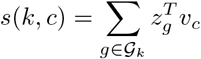

Where 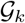 is the set of character n-grams of the k-mer *k*, *z* is a vector representation of all character n-grams and *v* is a k-mer vector. This method allows to learn a reliable representation even for rare k-mers. It can be useful since we don’t know the true size of each k-mer and where they are separated.

## D Multiple instance learning

*Zaheer* et al. (2017) defined a function invariant to permutation and place it in a neural network named *DeepSets*. The model is separated into 3 steps. A first neural network Phi projects each instance of the bags into a lower representation. A MIL layer (aggregation operation) invariant to permutation aggregates all instance in a bag. Finally, a last neural network operates on a single instance (aggregated by the preceding step) to compute prediction.

Mathematically *Zaheer* et al. (2017) defined a property and proved a theorem:

- Invariance to permutation can be formulate like this:

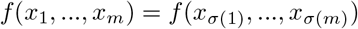

With *m* the number of elements in the bag and *σ* any permutation.
- Theorem: A function *S*(*X*) operating on a set *X* can be a valid scoring function i.e it is permutation invariant to the elements in X, if and only if it can be decomposed in the forme *ρ*(∑_*x*∈*X*_ *ϕ*(*x*))

The last theorem gives a structure for the neural network: *ϕ* the first neural network, ∑_*x*∈*X*_ the aggregation function and *ρ* the last neural network for classification. The sum operation is trivially invariant to permutation but we can also define other functions like mean or max pooling. Those aggregation functions are quite basic since they are not learned by the network. That’s why *Ilse* et al. (2018) have recently proposed a new method to parameterize all transformations. They defined a new function based on an attention mechanism (*Luong* et al. 2015). The proposed MIL pooling is:

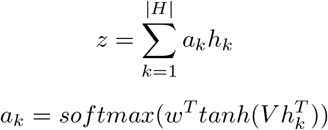

Where *w* ∈ ℝ^*n*^ and *V* ∈ ℝ^*n*×*m*^ are parameters and *H* = *h*_1_, ..., *h_k_* is a bag of *k* genomes embeddings. To be invariant to permutation the weights’ sum must be equal to one.

## E Code implementation

The full Pipeline is developed to train k-mers embeddings, reads embeddings, classfier of reads into taxa and classifier of metagenomes into disease. The algorithms and frameworks used for experiments are summarized below.

## Data processing

Given the amount of data we decided to take advantage of the Spark 2.4.5 python Framework (*Zaharia* et al. 2012) running on distributed clusters to manage memory and make parallel computing on CPU for all data processing. We use YARN or Torque as cluster managers to change the resources allocated for each of our experiments. The deep learning models are always trained on GPU computing.

## Machine learning

- *kmer2vec*: FastText, version 9.2 from GitHub written in C++.
- *read2vec*:

– FastText embeddings mean aggregation.
– Language model transformers, trained with Pytorch 1.6.
– FastDNA, from GitHub written in C++.
- *read2genome*:

– Multiple Layers Perceptron on top of read embeddings, H2O sparkling water 3.30.
– FastDNA, from GitHub written in C++.
- *metagenome2vec*:

– SVM, Random Forest, Gradient Boosting and Ada Boost from scikit-learn 0.22.1 for classification benchmark on *metagenome2vec* vectorial representation and *BoK* representation.
– DeepSets, using Pytorch 1.6, Ax 0.1.13 and ray-tune 0.8.5 packages for training and tuning the multiple instance learning model.

## F DNA embeddings intrinsic evaluation

## F.1 kmer2vec

K-mer embeddings are trained in a self-supervised manner that the algorithm tries to predict the surrounding k-mers in regard to the current one. The principal parameters that we can modify are the embeddings size (dimensionality complexity), the k-mer size (smaller or bigger pieces of DNA) and the window size (more or less surrounding words). It creates a big parameters space that influences severity points like the vocabulary size, the embeddings learning and more globally the final representation.

Analyzing k-mers vectors and finding best hyper parameters are done by intrinsic evaluation of the embeddings. The intrinsic evaluation is an important test that can help to identify if the algorithms learned good DNA embeddings. Unfortunately, this is not an obvious task according to DNA sequences. There is not a lot of information about the vocabulary compared to natural language where we assume that the vectors of two words like synonyms have a high similarity. Several intrinsic evaluation methods for NLP word embeddings are enumerated in *Bakarov* (2018) but none of them can be used with DNA because they rely on text-specific concepts. To overcome the fact that these methods are not available to evaluate the DNA embeddings, distance between k-mer chains is taken in account. *Ng* (2017) measure the relation between the cosine similarity of two vectors with their corresponding k-mers Needleman-Wunch score. In *Min* et al. (2017), authors prefer to compute the relation between the cosine similarity and the Edit distance. Both Edit distance and Needleman-Wunsch scores are computed on k-mer chains and compared to the cosine similarity of their embeddings. Figure 8a and 8b confirm that the distance between k-mers and between their embeddings do correlate. Unfortunately, these methods are only feasible when *k* is not too high, generally less than or equal to 6. Indeed, when *k* increases, so does the number of k-mers in vocabulary, which makes the calculation of distances much too long.

**Figure 8:**
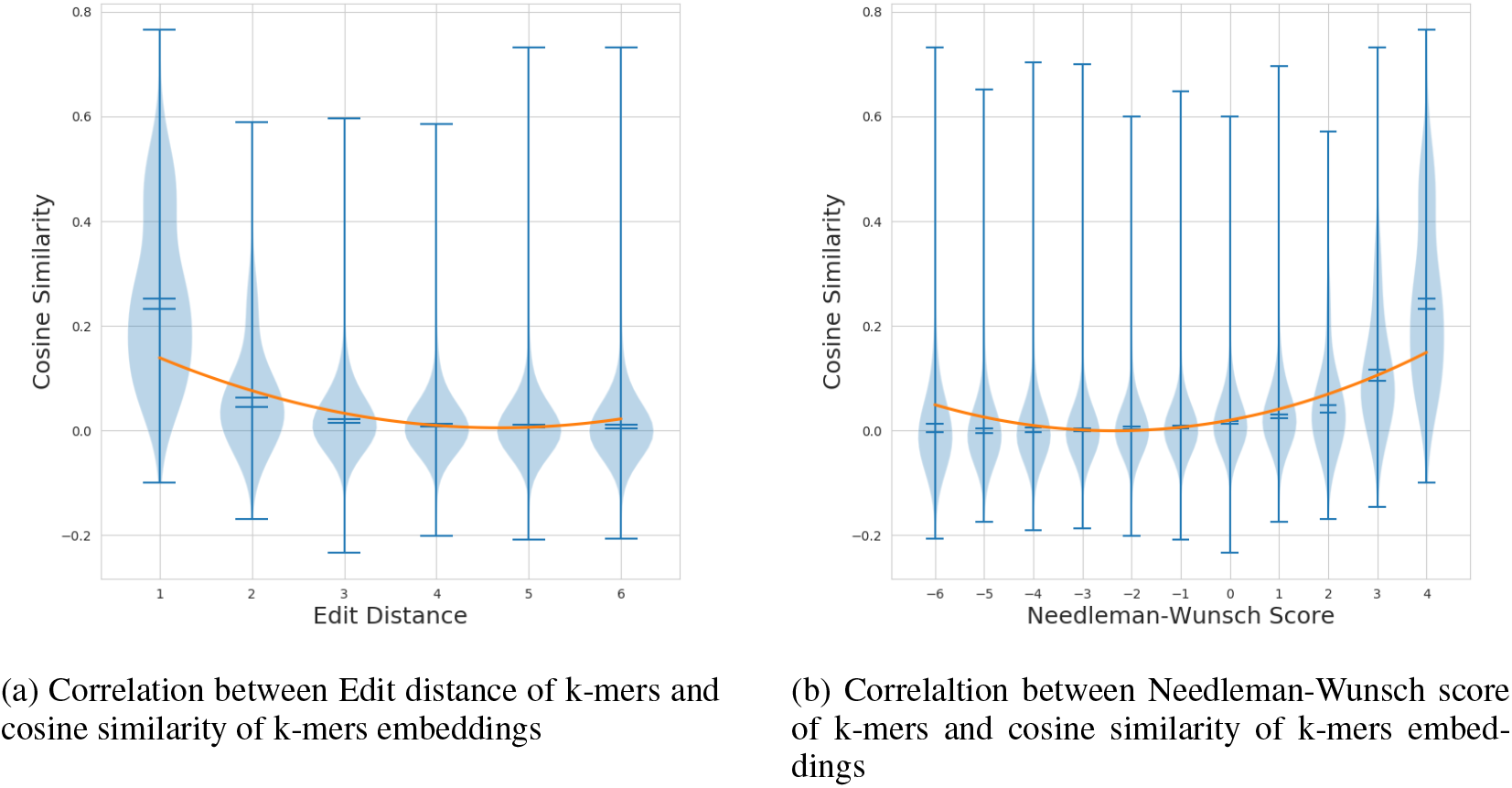
Each violin plot shows mean, median and the extreme values at a specific score. A smaller Edit distance and a higher Needleman-Wunsch score implies that k-mers are more similar. A higher cosine similarity implies that vectors are more co-linear.

## F.2 read2vec

Embeddings at the read level can’t benefit from analysis in Appendix F.1 because reads’ length are a lot bigger than k-mers. Nevertheless, as a genomes catalog has been used to train the read embeddings, it can be projected in this new vector space. Then, species from the same genus or with a similar genetic material should be more closely related to each other. We have set up two methods to observe this phenomenon. One is to project and visualize genomes embeddings using the t-SNE algorithm. Results on Figure 9 highlight that some clusters are formed of genomes from the same family. The other method is to compute a Mantel test and compare the correlation between two distance matrices of genomes. First is the cosine similarity between genome embeddings, second is the Mash distance which is a genome distance estimation using the MinHash algorithm between genome DNA. It is computed using the GitLab from the study of *Criscuolo* (2019). A high value in the mantel test implies that cosine similarity of the embeddings is correlated with the mash distance of DNA, then it gives a good indicator on the relevance of the representation learnt by the model. Models are tuned and results are reported in Table 3. *FastDNA* has the highest scores in this analysis. *Transformer* has better results than *FastText* for similar *k*. However it could not benefit from learning with a bigger *k* due to the complexity and the memory footprint of the model increasing exponentially with the size of the vocabulary. This is a blocking point because it is shown that increasing *k* leads to higher scores.

**Figure 9:**
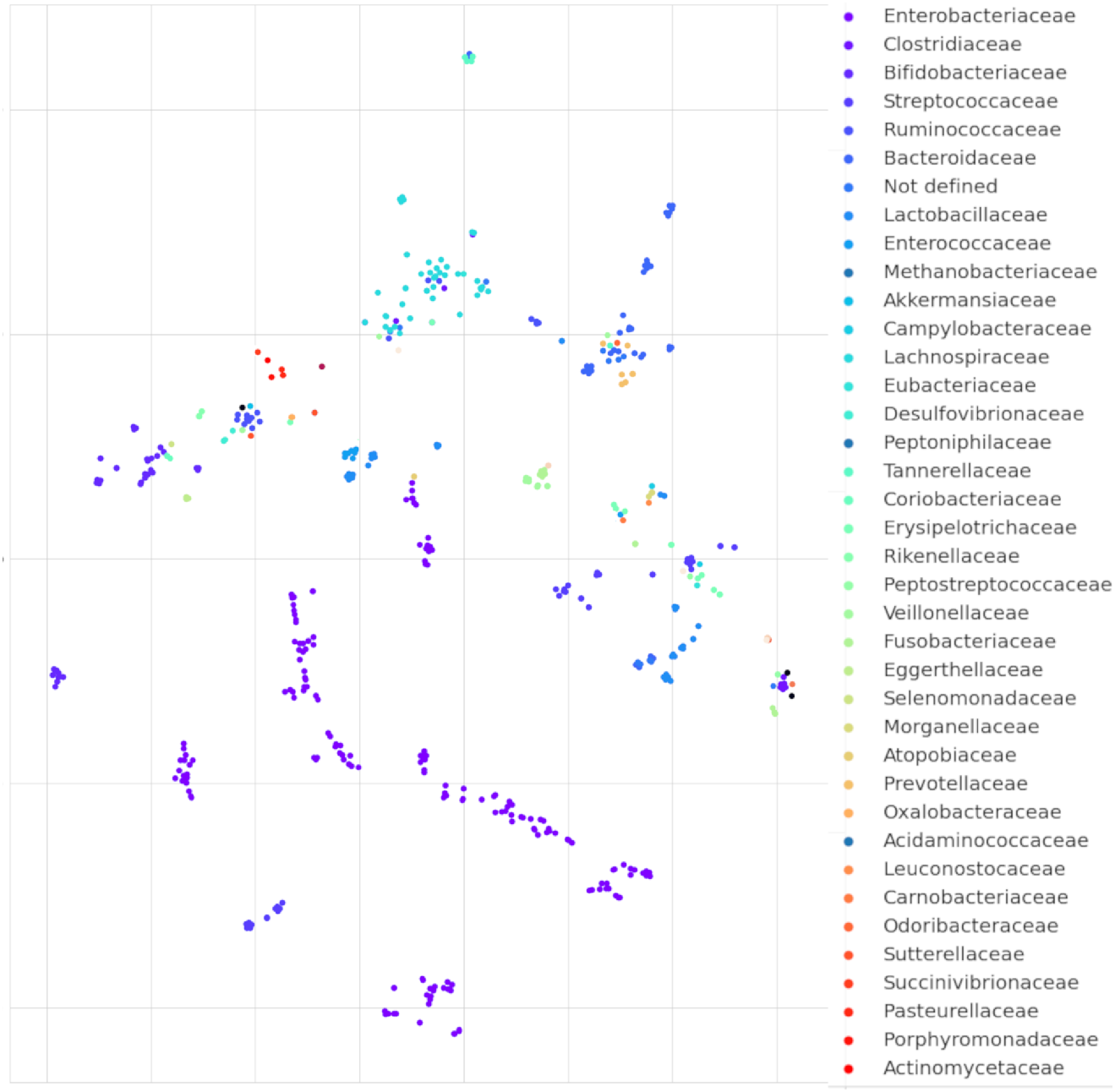
Each point is the projection into a 2D space with t-SNE algorithm of genome embeddings from FastDNA model. Points similarly colored have the same family.

**Table 3:** Mantel Test scores between Mash distance and genomes embeddings. Parameters *k* and *dim* refer to k-mer size and embeddings dimension respectively. When *k* increases the Mantel test gets a better value. FastDNA dominates the ranking in front of the *Transformer* and *FastText*.

| Method | Mantel Test |
| --- | --- |
| FastText k=3,dim=300 | 0.50 |
| FastText k=6,dim=300 | 0.54 |
| FastText k=9,dim=300 | 0.56 |
| FastText k=13,dim=100 | 0.61 |
| Transformer k=3,dim=300 | 0.52 |
| Transformer k=6,dim=300 | 0.56 |
| Transformer k=9,dim=300 | 0.59 |
| FastDNA k=13,dim=100 | 0.64 |

## G Read classification evaluation

We ensure that the *read2genome* model returns probabilities associated with the prediction, in that way we can set a threshold to reject classifications with too much uncertainty. As there is a tremendously high number of sequences by metagenome, rejecting uncertain predictions improves the precision of the model without impacting clusters of reads. Q. *Liang* et al. (2020) also uses a reject threshold determined manually in *DeepMicrobes*.

We compare the results of *FastDNA* and *transformer+MLP* models trained over 10 of the 235 species in the dataset. As *FastDNA* obtains the best scores on 10 species, we trained the model on the whole 235 species with parameters set to 13 for k-mer size, 100 for embedding dimension and 30 for the number of epochs. We compute and plot the accuracy, precision, recall, f2-score and rejected rate in accordance to the rejected threshold (see Figure 10). Metrics’ formula are recalled bellow:

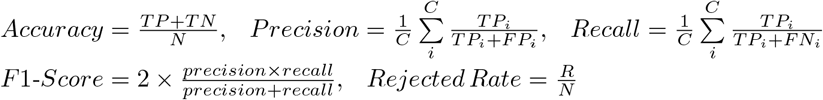

With *N* = # of samples, *C* = # of classes, *R* = # of rejected samples, *TP* = True Positive, *T N* = True Negative, *F P* = False Positive, *F N* = False Negative

**Figure 10:**
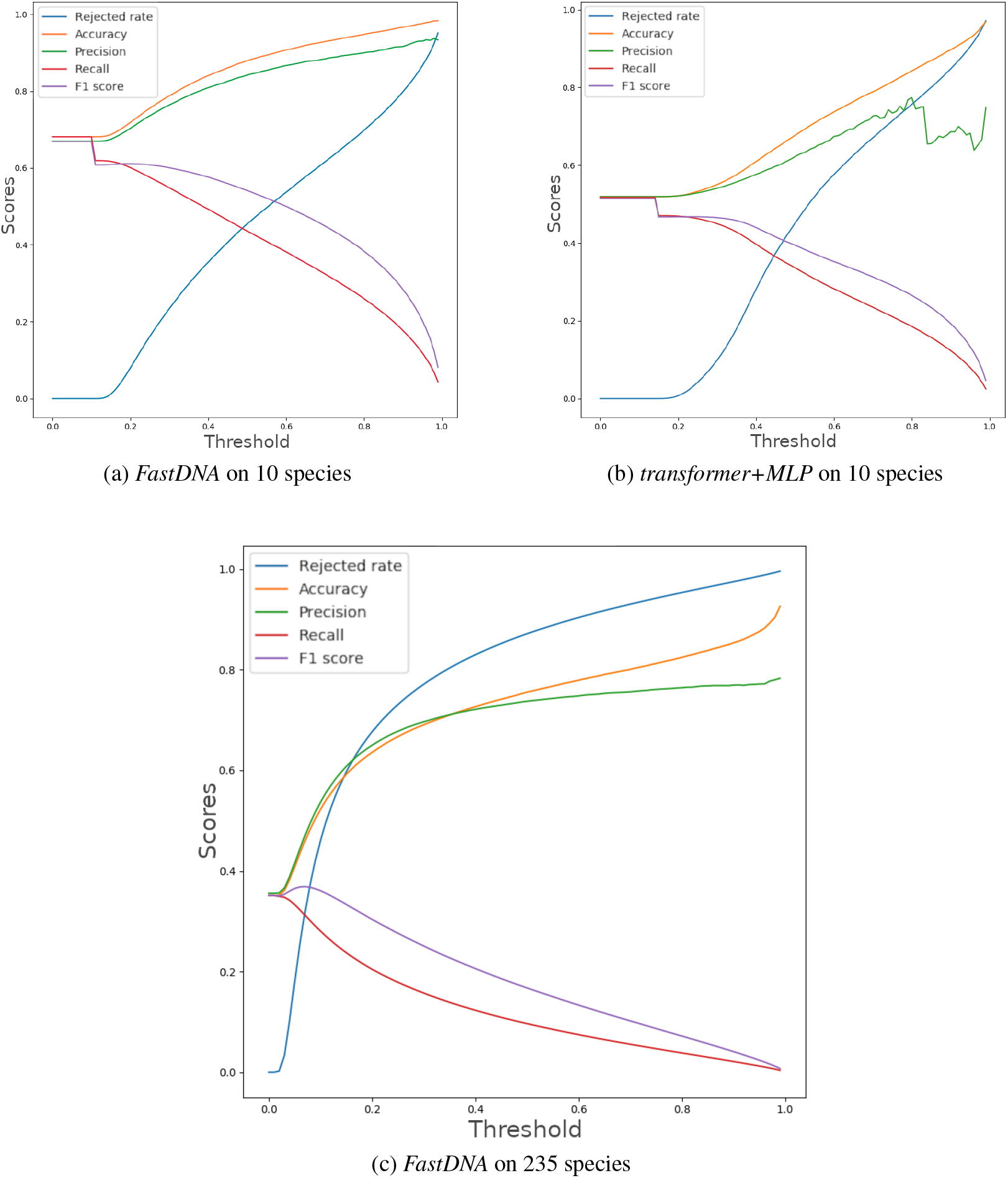
Scores obtained to classify reads into species from the 2.19M simulated reads of the validation dataset. The threshold corresponds to the minimum probability of the class predicted by the model so that the read is not rejected.

## H Metagenomes clustering

*m*_1_ metagenomes are selected and *m*_2_ < *m*_1_ others are cut in 10 sub parts. Each metagenome or part of the metagenome is represented by one vector. An agglomerative clustering is trained on these embeddings to compute a clustermap and show distances between them (Figure 11). Results show logically that embeddings from portions of the same metagenome are closer to each other. It also indicates this relation for metagenomes from the same class even if some of them are in the wrong cluster.

**Figure 11:**
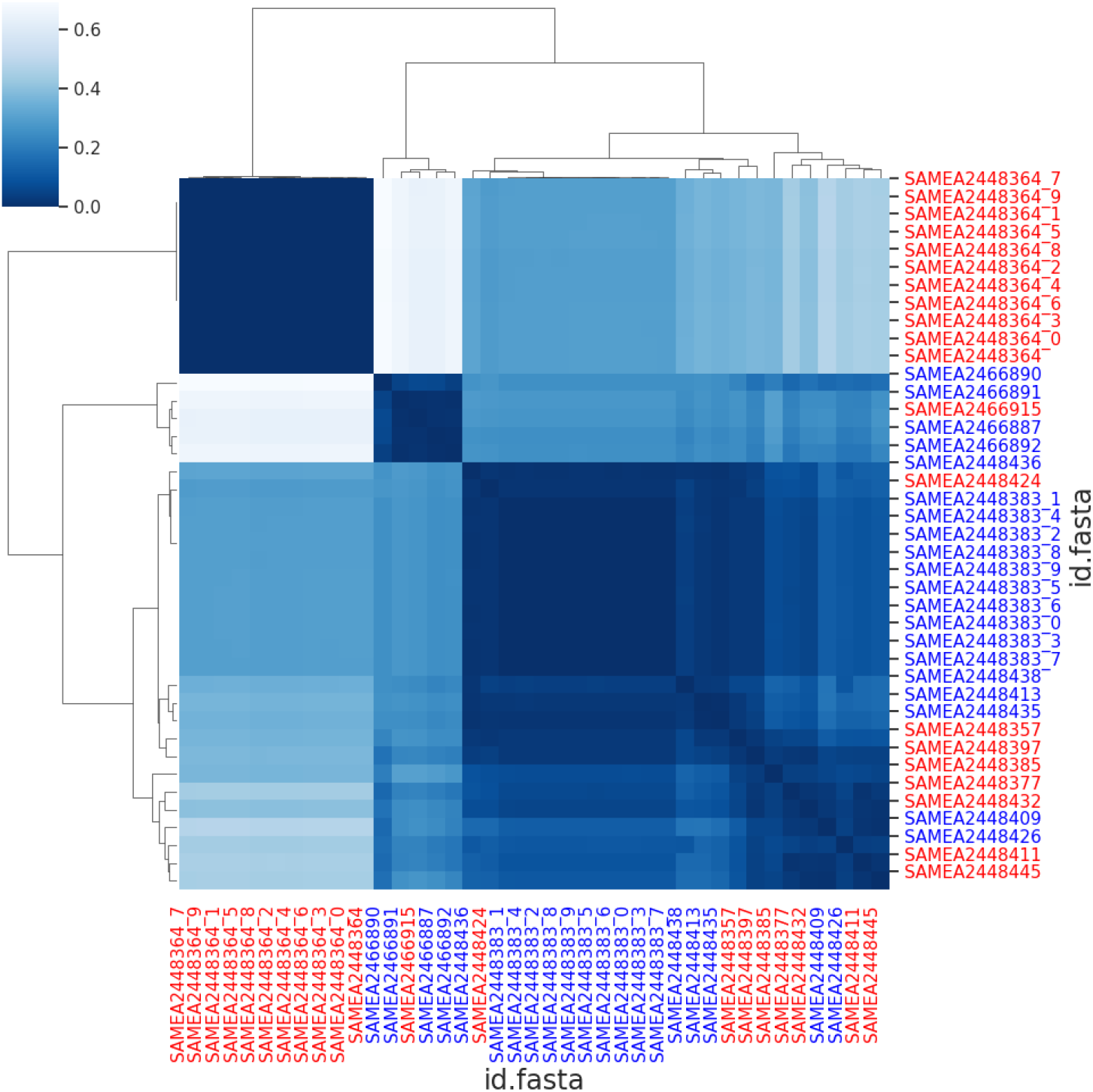
Cluster map computed on the Colorectal Dataset with the *metagenome2vec* vectorial representation. Blue ids and red ids refer to healthy patients and sick patients respectively. Underscores on ids followed by a digit correspond to partitions of a same metagenome. On the map, the darker the color, the more similar the metagenomes.

## Footnotes

1 Datasets have a massive size considering they only have few hundreds of patients. Metagenomes are composed of ~ 80 million reads, each one composed of ~ 90 nucleotides. It is a case where *D >> N* with *D* the number of features and *N* the number of examples. This is, in general, a complex task for machine learning algorithms.

## Notes

### Competing Interest Statement

The authors have declared no competing interest.

### Summary of Updates

Add author

